# Arabidopsis FIBRILLIN6 regulates carotenoid biosynthesis by directly promoting phytoene synthase activity

**DOI:** 10.1101/2022.06.30.498318

**Authors:** Ariadna Iglesias-Sanchez, Luca Morelli, Manuel Rodriguez-Concepcion

**Author notes:** Department of Biology & CESAM & ECOMARE, Universidade de Aveiro, 3810-193 Aveiro, Portugal (Luca Morelli).

## Abstract

Carotenoids are health-promoting plastidial isoprenoids with essential functions in plants as photoprotectants and photosynthetic pigments in chloroplasts. They also accumulate in specialized plastids named chromoplasts, providing color to non-photosynthetic tissues such as flower petals and ripe fruit. Carotenoid accumulation in chromoplast requires specialized structures and proteins such as fibrillins. Although fibrillins were first reported as structural components of carotenoid sequestering structures in chromoplasts, later work revealed roles in chloroplasts and other plastid types. However, the association of fibrillins with carotenoids in plastids other than chromoplasts has remained unexplored. Here we show that a member of the fibrillin family, FBN6, interacts with phytoene synthase (PSY, the first committed and rate-determining step of the carotenoid pathway) to promote its enzymatic activity. Transient overexpression of FBN6 in *Nicotiana benthamiana* leaves results in a higher production of phytoene, the product of PSY activity, whereas loss of FBN6 activity in *Arabidopsis thaliana* mutants dramatically reduces the production of carotenoids during seedling deetiolation and after exposure to high light. Our work hence demonstrates that fibrillins not only promote the accumulation of carotenoids but also their biosynthesis.

## Introduction

Carotenoids are a family of plant isoprenoids with roles in photosynthesis and photoprotection in green tissues. They also function as pigments in non-photosynthetic tissues such as flower petals and ripe fruits to attract animals for pollination and seed dispersal. Carotenoids are also precursors of bioactive molecules in plants (including hormones such as abscisic acid and strigolactones) and animals (including retinoids such as vitamin A) (Rodriguez-Concepcion et al., 2018). Carotenoids accumulate at high levels in chloroplasts, where they are mostly associated to the photosynthetic apparatus. However, they are most abundant in chromoplasts, which are plastids specialized in the sequestration of these highly lipophilic compounds found in carotenoid-accumulating non-photosynthetic tissues (Rodriguez-Concepcion et al., 2018). Chromoplast ultrastructure changes depending on the species, organ and carotenoid type they accumulate. For example, chromoplasts from carrot roots, daffodil petals and tomato ripe fruits accumulate carotenoids as large crystals whereas those of red pepper contain a large number of plastoglobules (PG) with fibrillar extensions surrounded by an outer layer of proteins referred to as plastoglobulins or fibrillins (Egea et al., 2010). Fibrillins (FBNs) also allow the sequestration of carotenoid (lycopene) crystals within membrane structures in tomato fruit during ripening.

FBNs are encoded by gene families in plants and algae that can be grouped in 12 clades (Singh and McNellis, 2011; Kim and Kim, 2022). They include members with diverse molecular masses, pI values, hydrophobicity profiles and homology with lipocalins (small proteins involved in the binding and transport of small lipophilic compounds), suggesting that each FBN family member has specific biological function(s) (Singh and McNellis, 2011; Lundquist et al., 2012; Kim and Kim, 2022). In agreement, plastid types vary in their FBN composition, suggesting that specific FBNs might have specialized functions in different classes of plastids (Singh and McNellis, 2011; Kim and Kim, 2022). However, their mechanisms of action remain poorly understood in most cases. In tomato, the PG-associated FBN1, FBN2 and FBN4 isoforms accumulate at high levels during fruit ripening, when carotenoids are actively produced and stored in chromoplasts (Barsan et al., 2012; Suzuki et al., 2015). By contrast, the *Arabidopsis thaliana* homologues FBN1a, FBN1b and FBN2 are involved in photoprotection and stress responses (Torres-Romero et al., 2022) whereas FBN4 is likely involved in the partitioning of metabolites between the PG and the thylakoid membranes of chloroplasts (Singh et al., 2012). FBN1a, FBN1b and FBN2 interact with each other around the PG surface, but FBN2 also plays FBN1-independent roles through interaction with other proteins (Torres-Romero et al., 2022). Arabidopsis FBN5 is a stromal protein involved in the acclimation to photooxidative stress but also in plastoquinone-9 biosynthesis by binding to solanesyl diphosphate synthases (Kim et al., 2015). FBN6 is needed for plants to acclimate to light stress and it might be part of the ROS scavenging machinery (Lee et al., 2020). The role of other FBNs remains poorly studied (Singh and McNellis, 2011; Kim and Kim, 2022).

FBN6 was found to be among the proteins co-immunoprecipitated with a GFP-tagged version of phytoene synthase (PSY), the first and main-rate-determining enzyme of the carotenoid pathway (Welsch et al., 2018). FBN6 was found to be the only FBN enriched in purified envelope fractions, where PSY is also found (Ferro et al., 2010; Bouchnak et al., 2019). We therefore wondered whether this particular isoform might modulate PSY activity, hence potentially impacting the metabolic flux of the carotenoid pathway. Besides confirming binding and co-localization of Arabidopsis FBN6 and PSY, here we demonstrate the functional relevance of such interaction.

## Results and Discussion

### Arabidopsis FBN6 and PSY physically interact in particular subplastidial locations

To confirm the previously observed interaction of Arabidopsis FBN6 and PSY proteins (Welsch et al., 2018) and test its specificity, we performed co-immunoprecipitation experiments in *Nicotiana benthamiana* leaves. Constructs encoding C-terminal myc-tagged FBN6 and GFP-tagged PSY were co-agroinfiltrated in leaves and 3 days later protein extracts were used for immunoprecipitation using anti-myc antibodies (Fig. 1). As a negative control, we used a myc-tagged version of Arabidopsis phosphoribulokinase (PRK), a stromal enzyme of the Calvin cycle (Barja et al., 2021). Immunoblot analysis of immunoprecipitated proteins with anti-GFP antibodies detected PSY-GFP in FBN6-myc but not in PRK-myc samples (Fig. 1A). The specificity of the FBN6-PSY interaction was then tested by using a myc-tagged version of the FBN4 protein as a negative control (Torres-Romero et al., 2022). The result (Fig. 1B) showed that Arabidopsis FBN4 does not interact with PSY and confirmed that interaction of FBN6 and PSY was specific.

**Figure 1.**
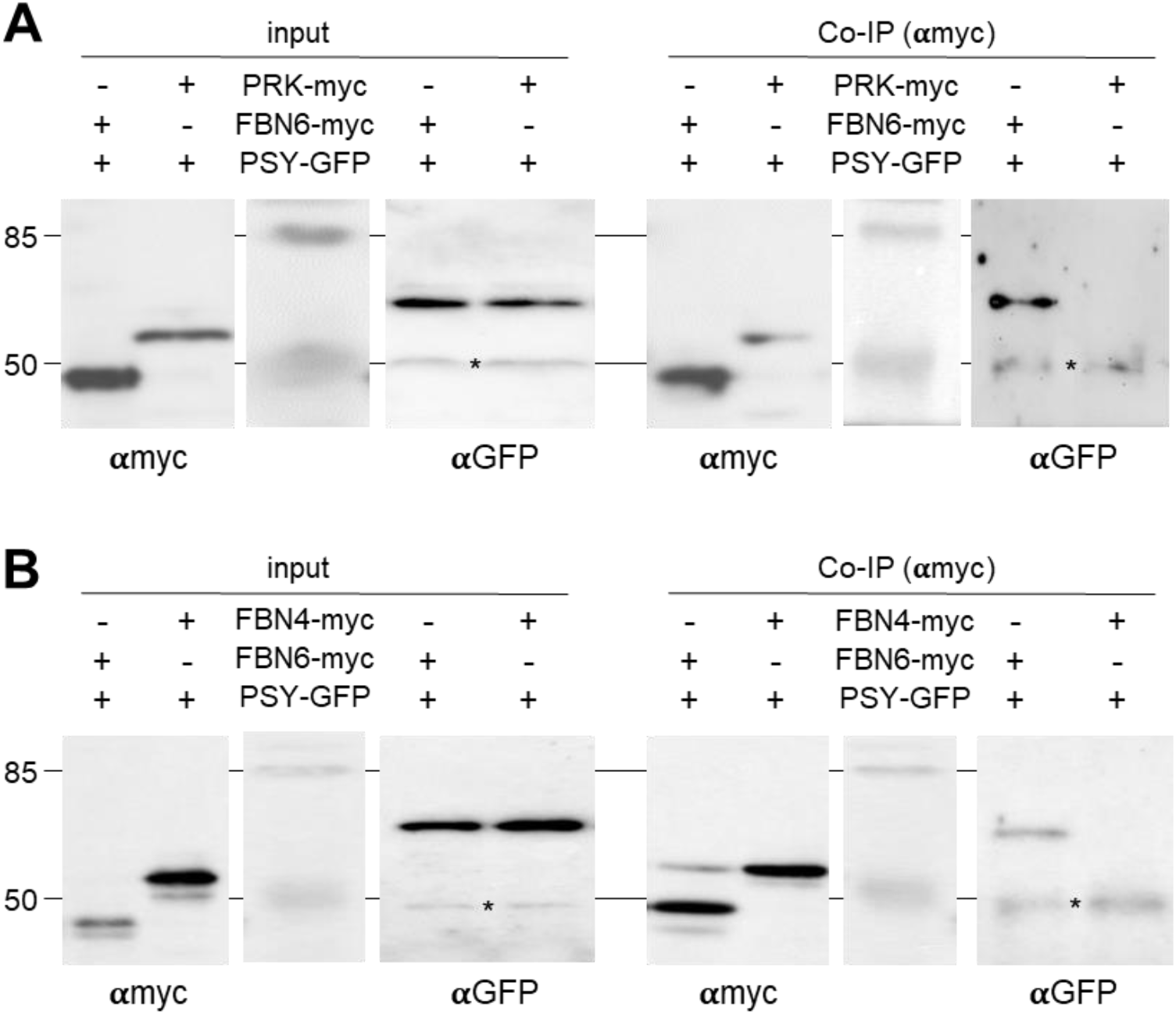
FBN6 and PSY can be immunoprecipitated together. *N. benthamiana* leaves were co-agroinfiltrated with the indicated combinations of proteins with C-terminal myc or GFP tags. Four days later, agroinfiltrated leaves were collected and used for protein extraction and analysis. Part of the protein extracts was used to test protein production (input) by immunoblot analyses with antibodies against myc (αmyc) or GFP (α-GFP; asterisk marks the position of unspecific bands). The remaining protein extracts were used for co-immunoprecipitation (Co-IP) experiments using αmyc followed by immunoblot analyses with immunoblot analyses with αmyc (to confirm successful IP) and αGFP (to detect the presence of co-immunoprecipitated PSY-GFP protein). (A) Experiment using the Calvin cycle enzyme PRK fused to the myc tag as a negative control. (B) Experiment using the PG-associated FBN4 protein fused to the myc tag to test for the specificity of the FBN6-PSY interaction.

When analyzing the co-immunoprecipitation assays, we noticed that the PSY-GFP protein concentrated in fluorescent dots within chloroplasts (Fig. 2A). These dots are similar to those observed for PG-associated proteins (Lundquist et al., 2012; Gámez-Arjona et al., 2014; Morelli et al., 2022) and also for PSY enzymes (Shumskaya et al., 2012) fused to fluorescent protein tags. Interestingly, GFP-tagged Arabidopsis FBN6 also shows a spotted distribution within chloroplasts despite the protein is not found in PG but localized in thylakoid and envelope membranes (Ferro et al., 2010; Lundquist et al., 2012; Lee et al., 2020; Bouchnak et al., 2019). To test whether the fluorescent dots observed in chloroplasts harboring FBN6-GFP and PSY-GFP proteins corresponded to the same or different subplastidial locations, we generated a FBN6-RFP construct and co-agroinfiltrated it with the PSY-GFP construct. Confocal microscopy showed a clear overlapping of RFP and GFP fluorescence signals (Fig. 2A), indicating that FBN6 and PSY co-localize within chloroplasts, as expected for interacting proteins. Despite FBN6-GFP has been previously found to co-localize with PG proteins fused to fluorescent tags in co-expression experiments very similar to those reported here, subsequent analysis of chloroplast membrane fractions argued against a localization of FBN6 in PG (Lee et al., 2020), a conclusion that is also supported by proteomic data (Ferro et al., 2010; Lundquist et al., 2012; Bouchnak et al., 2019). While overexpression of FBNs and other PG-associated proteins typically results in PG proliferation (Shanmugabalaji et al., 2013; Van Wijk and Kessler, 2017), overproduction of FBN6 or/and PSY proteins in agroinfiltrated leaf chloroplasts did not cause changes in PG core-associated proteins such as FBN1 or FBN2 homologues (Fig. 2B). These results provide further indirect evidence that FBN6 and PSY are not PG-associated proteins. Together, we conclude that Arabidopsis FBN6 and PSY interact and preferentially co-localize in particular locations of the chloroplast such as envelope or thylakoid membrane domains different from PG.

**Figure 2.**
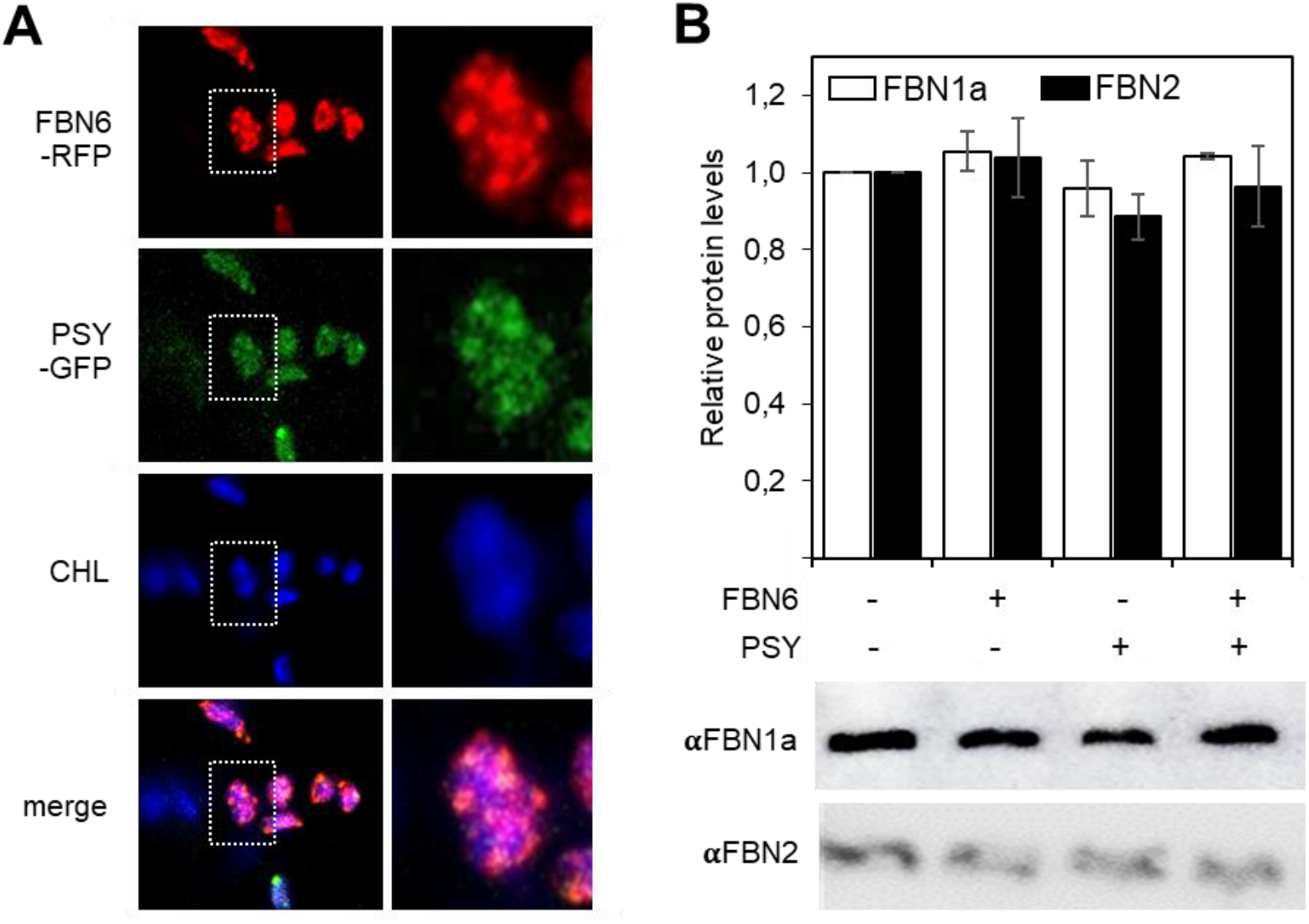
FBN6 and PSY co-localize in chloroplasts. *N. benthamiana* leaves were co-agroinfiltrated with constructs encoding FBN6-RFP and PSY-GFP and four days later samples were taken for confocal and immunoblot analyses. (A) Representative confocal microscopy images of a leaf section transiently expressing the indicated fluorescent fusion proteins. RFP fluorescence is shown in red, GFP fluorescence in green, and chlorophyll (CHL) autofluorescence in blue. Bottom panel shows a merge of the three fluorescence channels. Right panels are a magnification of the area boxed in white in the left panels. (B) Immunoblot analysis of the accumulation of FBNs identified using antibodies against Arabidopsis PG-associated FBN1a and FBN2 proteins in leaves expressing the indicated constructs or a GFP control (-/-). Quantification was made by relativizing each band intensity to the one obtained in the control. Representative results are shown in the lower panels. Barplot shows the mean ± SD values of quantitative data from n=3 leaves.

### FBN6 promotes PSY activity

To test the possible relevance of FBN6 for PSY activity, we analyzed the production of phytoene (the direct product of PSY activity) and downstream carotenoids in *N. benthamiana* leaves agroinfiltrated with construct to transiently express the Arabidopsis genes encoding FBN6, PSY, or both (Fig. 3). Control leaves were agroinfiltrated with a GFP construct. While phytoene was not detected in control or FBN6 samples, it was present in PSY samples and it increased in samples agroinfiltrated with both FBN6 and PSY (Fig. 3A). Treatment of agroinfiltrated leaves with norflurazon (NF), an inhibitor that prevents phytoene conversion into downstream carotenoids, allowed to detect phytoene in all the samples. Most interestingly, FBN6 samples showed higher levels of phytoene than GFP controls, supporting the conclusion that FBN6 is a promoter of the endogenous PSY activity. The presence FBN6 also led to highly increased phytoene levels in *PSY*-overexpressing samples (Fig. 3A). In the case of total carotenoids, FBN6 alone did not lead to increased levels but a combination of FBN6 and PSY resulted in more carotenoids that PSY alone (Fig. 3B). These results suggest that PSY-derived phytoene limits the production of downstream carotenoids in leaves and confirm a positive effect of FBN6 on PSY activity.

**Figure 3.**
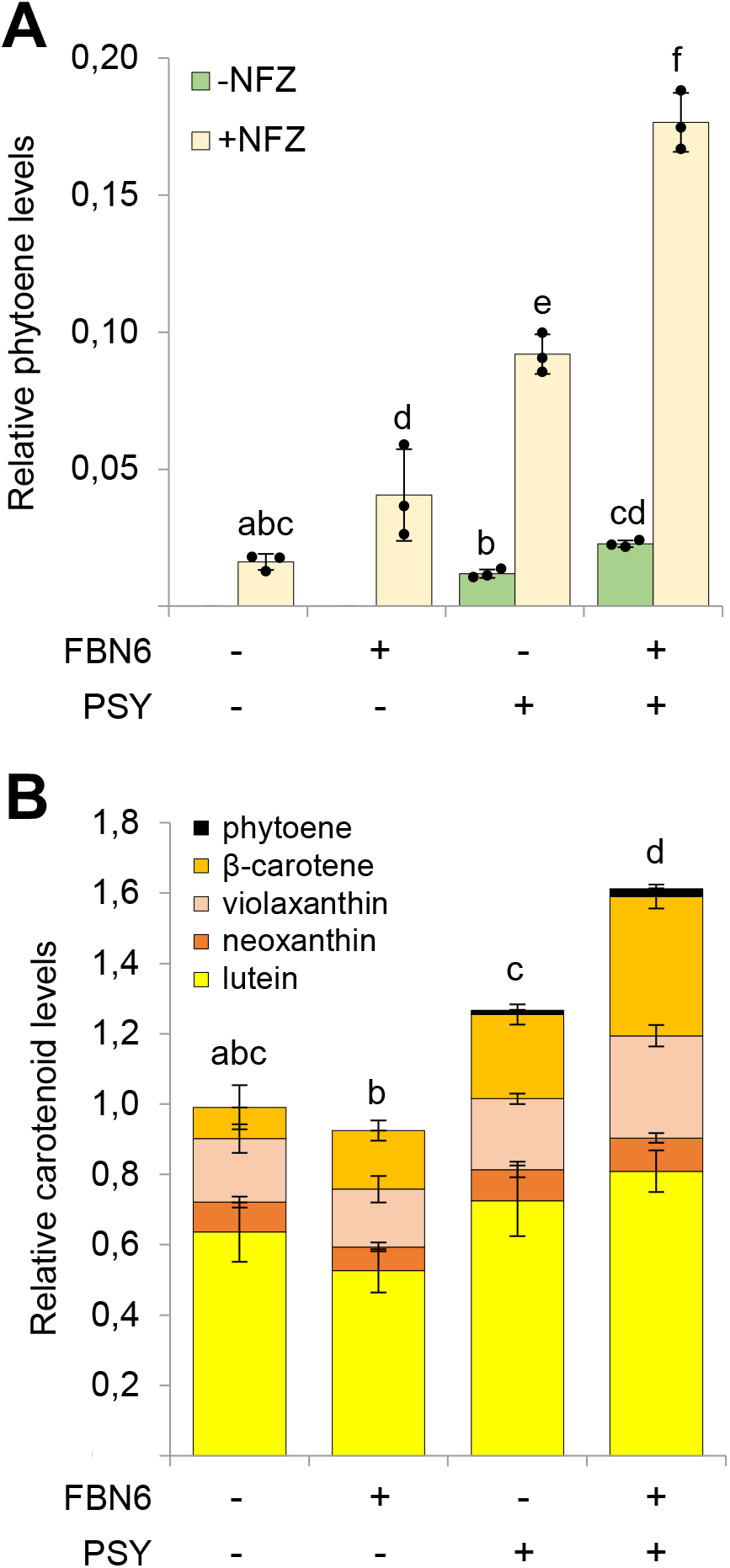
FBN6 promotes phytoene and downstream carotenoid production. *N. benthamiana* leaves were co-agroinfiltrated with constructs encoding FBN6 or/and PSY and 4 days later samples were taken for HPLC analysis of carotenoids. Control experiments (-/-) were carried out with a GFP construct. (A) Phytoene levels in leaf samples either treated (+) or not (-) with norflurazon (NF). Values are represented relative to total carotenoid levels in untreated control leaves and they correspond to the mean ± SD of n=3 leaf samples. (B) Carotenoid levels in – NF samples. Values are represented relative to total carotenoid levels in control leaves and they correspond to the mean ± SD of n=3 leaf samples. In both plots, different letters represent statistically significant differences (*p* < 0.05) among means according to one-way ANOVA followed by post hoc Tukey’s tests.

If FBN6 is a positive regulator of PSY, loss of FBN6 in mutants would be expected to prevent normal carotenoid synthesis. FBN6-defective Arabidopsis *fbn6-1* mutants were previously reported to have reduced chlorophyll levels compared to wild-type (WT) controls, but no data on carotenoid levels were reported (Lee et al., 2020). In Arabidopsis, the most active production of carotenoids takes place during deetiolation to protect the emerging photosynthetic apparatus against photooxidative damage caused by excess incoming light (Rodríguez-Villalón et al., 2009). The boost in carotenoid synthesis during deetiolation is mostly mediated by increased expression of the PSY-encoding gene in Arabidopsis (Rodríguez-Villalón et al., 2009; Toledo-Ortiz et al., 2010). Interestingly, the gene encoding FBN6 was also highly induced during this process (Fig. 4A). Compared to the WT, the *fbn6-1* mutant accumulates slightly less carotenoids in etiolated seedlings and shows a strongly attenuated production during deetiolation (up to 24h after illumination). As a result, greening (chlorophyll accumulation) was delayed (Fig. 4B). After a few days under normal light (NL), the differences in carotenoid (and chlorophyll) levels between WT and *fbn6-1* plants are attenuated but still detectable (Fig. 4C).

**Figure 4.**
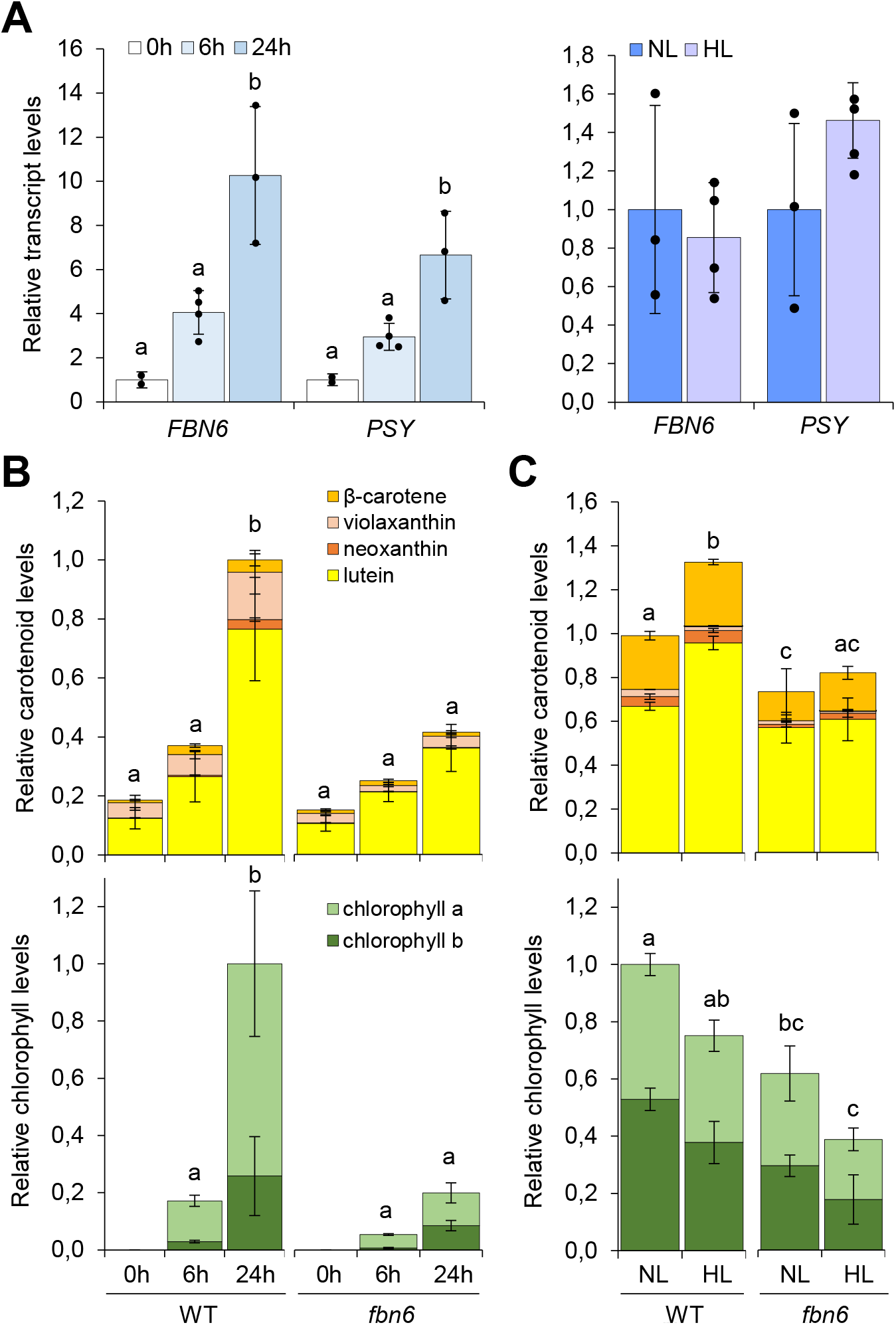
FBN6 is required for carotenoid production in Arabidopsis. Deetiolation experiments were carried out with Arabidopsis WT and FBN6-defective mutant seeds germinated in the dark for 3 days; the resulting etiolated seedlings were then exposed to white light for 6h and 24h. For high-light (HL) experiments, WT and *fbn6* seedlings germinated and grown under normal light (NL) conditions for 9 days were exposed for 24h to either NL (as a control) or HL. (A) Expression profiles of genes encoding FBN6 and PSY during deetiolation (left) or HL exposure (right) of WT seedlings. Transcript levels are normalized using the *UBC21* gene and are shown relative to etiolated (0h) or NL-exposed samples, respectively. (B) Carotenoid and chlorophyll levels in deetiolating WT and *fbn6* seedlings. Values are represented relative to total pigment levels in WT 24h samples. (C) Carotenoid and chlorophyll levels in HL-exposed WT and *fbn6* seedlings. Values are represented relative to total pigment levels in WT NL samples. In all plots, mean ± SD of n≥3 replicates are represented. Letters represent statistically significant differences (*p* < 0.05) among means according to one-way ANOVA followed by post hoc Tukey’s tests.

Carotenoid levels, but not *PSY* expression, increase when Arabidopsis plants are exposed to high light (HL), suggesting an up-regulation of PSY enzyme activity to increase the metabolic flux towards carotenoid synthesis (Kreslavski et al., 2021). Similar to *PSY*, transcripts encoding FBN6 also showed unchanged levels when 9-day-old Arabidopsis seedlings were exposed for 24h to HL compared to controls left at NL for the same time (Fig. 4A). Most interestingly, the HL-triggered increase in carotenoid levels was highly attenuated in FBN6-defective plants compared to the WT controls (Fig. 4C). By contrast, the reduction in chlorophylls associated to HL exposure was similar in WT and *fbn6-1* seedlings in absolute terms but seemingly stronger in the mutant in relative terms (Fig. 4C), likely because disrupted up-regulation of carotenoid contents in *fbn6-1* seedlings results in defective photoprotection and a higher chlorophyll degradation rate. We conclude that FBN6 is required to support the increase in carotenoid levels that takes place in response to HL, most likely because it directly promotes PSY enzyme activity during this process.

### A new chapter in the connection between FBNs and carotenoids

FBN gene expression often correlates with the carotenoid content of non-photosynthetic plant tissues, e.g. during fruit ripening, where their role is assumed to be mainly structural (Singh and McNellis, 2011). Indeed, FBNs were first identified as proteins specifically associated to carotenoid accumulation in fruit and petal chromoplasts, as components of “fibrils” and PG structures (Deruère et al., 1994; Vainstein et al., 1994). Similarly, green algae FBNs were also proposed to be involved in the formation and stabilization of the chloroplast eyespot, a carotenoid-rich light sensing system (Schmidt et al., 2006; Davidi et al., 2015). In plant chloroplasts, however, carotenoids are not normally concentrated in PG or any other lipid body but distributed in thylakoids and, to a lower extent, envelope membranes (Morelli et al., 2022). Despite chloroplast FBNs are presumed to maintain photosynthetic function and they are present in all the compartments where carotenoids are synthesized and accumulated, a direct functional association between FBNs and carotenoid biosynthesis has remained unexplored.

Studies on FBN-interacting proteins have substantially improved our understanding of the function of different FBN isoforms. Arabidopsis FBN1a, FBN1b, and FBN2 interact with each other, presumably forming a network around the PG surface that might be important to recruit other FBN-interacting proteins to PG (Gámez-Arjona et al., 2014; Torres-Romero et al., 2022). For example, these FBNs interact with allene oxide synthase (AOS), which catalyzes one of the first steps of jasmonic acid (JA) biosynthesis. FBN2 also interacts with other proteins involved in different metabolic processes, including other enzymes of the JA pathway and the carotenoid cleavage dioxygenase CCD4, a PG-associated enzyme involved in carotenoid degradation (Torres-Romero et al., 2022). It was proposed that FBN2 might mediate the recruitment of JA biosynthetic enzymes to the PG under stress conditions, but the functional relevance of the interaction with CCD4 was not explored (Torres-Romero et al., 2022). FBN1a and FBN1b also interact with starch synthase isoform 4 (SS4), but the elimination of the FBNs does not alter the initiation of starch granule formation (Gámez-Arjona et al., 2014). FBN4 interacts with the major ferredoxin protein in Arabidopsis (Fd2) and with harpin protein HrpN, an elicitor secreted by pathogenic bacteria that activate plant defense and resistance (Kim and Kim, 2022). FBN5 participates in the production of the plastidial isoprenoid plastoquinone through interaction with solanesyl diphosphate synthase isoforms 1 and 2 (SPS1 and SPS2), which catalyze the first committed step in the production of the prenyl chain of the molecule (Kim et al., 2015). Based on the presence of a lipocalin domain involved in binding to and transport of small hydrophobic metabolites, it was proposed that FBN5 might bind the solanesyl diphosphate product of SPS1 and SPS2 and remove it from the active site of the enzymes, hence contributing to increasing their enzymatic activity (Kim et al., 2015).

The lipocalin motif of FBN4 likely contributes to the role of this isoform in the trafficking of plastoquinone and other small lipids between the thylakoids and the PG (Singh et al., 2012). Interestingly, FBN6 also contains a lipocalin motif (Singh and McNellis, 2011). Based on that proposed for FBN5, a non-PG FBN protein, we speculate that direct interaction of FBN6 with PSY might result in improved activity of the enzyme by facilitating removal of the phytoene product from the active site. Regardless of the specific mechanism involved, our data support the conclusion that FBN6 plays a physiologically relevant role for carotenoid biosynthesis in Arabidopsis chloroplasts by physically binding to PSY to promote its enzymatic activity. Therefore, we demonstrate that FBNs not only promote the accumulation of carotenoids but also their biosynthesis. The unveiled role of FBN6 in carotenogenesis reported here can explain the phenotypes previously reported for the *fbn6-1* mutant, which were similar to those of mutants disrupted in ROS homeostasis (Lee et al., 2020). Because carotenoids are essential photoprotectants that dissipate excess light energy as heat and detoxify excess ROS in chloroplasts (Rodriguez-Concepcion et al., 2018), we conclude that the reduced acclimation to light stress and impaired ROS scavenging phenotypes observed in the *fbn6-1* mutant are likely a primary result of defective carotenoid biosynthesis. Lower growth, increased glutathione levels or resistance to cadmium stress would be secondary effects of impaired ROS homeostasis.

Similar to the up-regulation of *PSY* and *FBN6* genes during Arabidopsis deetiolation (Fig. 4A), the burst in carotenoid synthesis that takes place during tomato fruit ripening is also associated with increased expression of genes encoding the corresponding PSY and FBN6 homologues (Barja et al., 2021; Sun et al., 2022). This observation suggests that FBN6-dependent promotion of PSY activity and carotenoid biosynthesis might not be restricted to chloroplasts but also occur in chromoplasts. Further experiments with FBN6-defective tomato plants should confirm this possibility and provide new insights on the possible biotechnological use of this FBN isoform.

## Methods

### Plant material and growth conditions

*Nicotiana benthamiana* and *Arabidopsis thaliana* plants were grown as described (Morelli et al., 2022). The *fbn6-1* mutant (GABI_159E10) (Lee et al., 2020) was obtained from the Nottingham Arabidopsis Stock Centre (NASC). Homozygous lines were identified by genomic PCR with T-DNA and gene-specific primers (Supplemental Table S1). For the deetiolation experiment, seeds were surface-sterilized and sown on Petri dishes containing solid 0.5X MS medium without sucrose. After 3 days at 4ºC, they were exposed to white light (50 μmol photons·m^−2^·s^−1^, referred to as normal light or NL) for 3h and then incubated at 22ºC in the dark for 3 more days before exposure to NL to promote deetiolation. For the intense light experiments, seedlings grown on plates for 9 days under NL were exposed for 24h to 600 μmol photons·m^−2^·s^−1^ (referred to as high light or HL) or left under NL for the same time. *Nicotiana benthamiana* leaves were agroinfiltrated as described (Morelli et al., 2022).

### Gene constructs

The full coding region of the Arabidopsis FBN6 (AT5G19940) cDNA was PCR-amplified using primers AtFBN6-attB1-F and AtFBN6-attB2-R (Supplemental Table S1) and cloned into Gateway pDONR207 and subsequently pGWB420 (to create the FBN6-myc fusion) and pGWB454 (to create the FBN6-RFP fusion). Similarly, the FBN4-myc fusion was generated by PCR-mediated amplification of a cDNA sequence encoding full-length FBN4 (AT3G23400) with primers AtFBN4-attB1-F and AtFBN4-attB2-R (Supplemental Table S1) and cloned into Gateway pGWB420 via pDONR207. Constructs encoding PRK-myc (Barja et al., 2021) and PSY-GFP (Welsch et al., 2018) were previously available in the lab.

### Microscopy

Subcellular localization of fluorescent proteins transiently expressed in agroinfiltrated *N. benthamiana* leaves was analyzed with a Leica TCS SP5 Confocal Laser Scanning Microscope at 4 days post-infiltration. GFP fluorescence was detected using a BP515-525 filter after excitation at 488 nm while RFP signal was detected at 588 nm after excitation at 532 nm. Chlorophyll autofluorescence was detected using a LP590 filter after excitation at 568 nm.

### Co-immunoprecipitation and immunoblot assays

*N. benthamiana* leaves were agroinfiltrated with the appropriate constructs and samples were collected after 4 days for co-IP experiments performed as described (Barja et al., 2021). Immunoblot analysis with antibodies against FBN1a and FBN2 and subsequent quantification of the signals was carried out as described (Morelli et al., 2022).

### Gene expression and metabolite analysis

RNA isolation, cDNA synthesis, and RT-qPCR experiments were carried out as described (Barja et al., 2021) using the primers listed in Supplemental Table S1. Normalized transcript levels were calculated using Arabidopsis *UBC21* (At5g25760) as the reference gene. Carotenoids and chlorophylls were extracted, separated and quantified by HPLC-DAD as described (Barja et al., 2021).

## Acknowledgments

We thank Venkatasalam Shanmugabalaji and Felix Kessler (University of Neuchatel, Switzerland) for the kind gift of the fibrillin antibodies, Tsuyoshi Nakagawa (Shimane University, Japan) for the Gateway vectors, and the Nottingham Arabidopsis Stock Centre (NASC) for providing *fbn6-1* seeds. We also thank members of our laboratory for helpful discussions. No conflict of interest is declared.

## Author contributions

AI-S, LM and MR-C designed the research and analyzed data; AI-S and LM conducted the experiments; MR-C wrote the paper.

## Supplemental data

**Supplemental Table S1. Primers used in this work**.

